# An Ensemble Metabolome-Epigenome Interaction Network Identifies Metabolite Modulators of Epigenetic Drugs

**DOI:** 10.1101/2023.02.27.530260

**Authors:** Scott E. Campit, Rupa Bhowmick, Taoan Lu, Aaditi Vivek Saoji, Ran Jin, Madeline R. Shay, Aaron M. Robida, Sriram Chandrasekaran

**Affiliations:** Program in Chemical Biology, University of Michigan, Ann Arbor; Department of Biomedical Engineering, University of Michigan, Ann Arbor; Department of Electrical Engineering and Computer Science, University of Michigan, Ann Arbor; Department of Cellular and Molecular Biology, University of Michigan, Ann Arbor; Program in Cellular and Molecular Biology, University of Michigan, Ann Arbor; Center for Chemical Genomics, University of Michigan, Ann Arbor; Rogel Cancer Center, University of Michigan Medical School, Ann Arbor, MI, 48109, USA

## Abstract

Metabolites such as acetyl-CoA and citrate play an important moonlighting role by influencing the levels of histone post-translational modifications (PTMs) and regulating gene expression. This cross talk between metabolism and epigenome impacts numerous biological processes including development and tumorigenesis. However, the extent of moonlighting activities of cellular metabolites in modulating the epigenome is unknown. We developed a data-driven screen to discover moonlighting metabolites by constructing a histone PTM-metabolite interaction network using global chromatin profiles, metabolomics, and epigenetic drug sensitivity data from over 600 cell lines. Our ensemble statistical learning approach uncovered metabolites that are predictive of histone PTM levels and epigenetic drug sensitivity. We experimentally validated synergistic and antagonistic interactions between histone deacetylase and demethylase inhibitors with epigenetic metabolites kynurenic acid, pantothenate, and 1-methylnicotinamide. We apply our approach to track metaboloepigenetic interactions during the epithelial-mesenchymal transition. Overall, our data-driven approach unveils a broader range of metaboloepigenetic interactions than anticipated from previous studies, with implications for reversing aberrant epigenetic alterations and enhancing epigenetic therapies through diet.

## Introduction

While the regulation of metabolism through transcriptional and signaling pathways is extensively studied, how metabolites feedback and regulate gene expression is poorly understood^1^. Several metabolites are known to regulate gene expression in eukaryotes via histone post-translational modifications (PTMs)^2,3^. Epigenetic enzymes that “write” or “erase” histone PTMs sense levels of specific metabolites, such as S-adenosylmethionine, acetyl-CoA, and NAD+^2–4^. Acetyl-CoA is the substrate for histone acetyltransferases (HATs)^5,6^, while sirtuins, a family of histone deacetylases (HDACs), remove acetyl and other acyl moieties from lysine residues using NAD+^7,8^. Histone methyltransferases (HMTs) utilize S-adenyosylmethionine (SAM) to install methyl groups on histone tails^9,10^ and multiple methyl groups (up to 3) can be installed to exert different expression patterns^11,12^. Histone demethylases remove methyl groups using FAD with alpha-ketoglutarate, or another mechanism that utilizes oxygen, iron, and L-ascorbate^11,12^.

Metabolic-epigenetic interactions play a central role in controlling cellular development and physiology. For instance, the activity of the one-carbon pathway impacts histone methylation levels, and acetyl-CoA levels correlate with histone acetylation in embryonic stem cells^2,5,13,14^. Metabolic dysregulation can lead to altered acetylation states associated with increased cancer risk^15^. Glucose-derived acetyl-CoA correlates with histone acetylation levels and activation of oncogenes in various cancer cell models^16,17^. Loss of function mutations in metabolic genes causes an accumulation of succinate, fumarate, and 2-hydroxyglutarate in diverse cancers. These metabolites inhibit demethylase enzymes such as the tumor suppressor TET2 and promote the epithelial-to-mesenchymal transition^2,18,19^. While we understand how genetic and signaling mechanisms influence metabolic dysregulation in cancer and other diseases^20–22^, the link between metabolism and gene expression via epigenetic regulation, termed as metaboloepigenetics^23^, is an active area of research and may benefit from a systems approach^1,24–27^.

Epigenome-targeted therapies are emerging as a therapeutic option for several cancers and immune disorders, with numerous drugs currently in the preclinical and clinical pipeline^28,29^. Vorinostat was the first HDAC inhibitor approved by the FDA to treat cutaneous T-cell lymphoma^30^. Since then, other HDAC inhibitors have been approved as anticancer agents. Additionally, several promising preclinical compounds inhibit histone methyltransferases and lysine demethylases for cancer treatment^28,31^.

Cellular metabolic status can impact the efficacy of drugs, including HDAC inhibitors^29^. Notably, flux balance analysis showed that the metabolic flux of acetyl-CoA is predictive of sensitivity to HDAC inhibitors^32^. Prior studies based on transcriptomics data have found that metabolic gene expression is predictive of anti-cancer drug sensitivity^33^. However, it is unknown how effective epigenetic therapeutics are in the context of environmental variables such as metabolite or nutrient levels. Thus, there is a need to develop tools that can predict the impact metabolism has on chromatin-modifying drugs. Further, the metabolic dependencies of epigenetic drugs can shed light on mechanistic interactions between metabolism and epigenome.

Here we apply a data-driven, systems-pharmacology approach^34,35^ to discover metaboloepigenetic interactions. We constructed an interaction network using global chromatin profiles, epigenetic drug sensitivity data, and metabolomics data from over 600 cancer cell lines in the Cancer Cell Line Encyclopedia (CCLE). Using an ensemble of machine learning models, we prioritized top histone PTM-metabolite and metabolite-drug interaction pairs. Our analysis uncovered surprising metabolic dependencies of epigenetic drugs, and we validated predictions for four metabolites that had synergistic/antagonistic relationships with chromatin-modifying drugs Vorinostat and GSK-J4. We further apply our interaction network to study metabolite/PTM interaction dynamics during the epithelial-mesenchymal transition (EMT). Together, our findings provide a comprehensive resource to explore the complex relationship between cellular metabolism, epigenetics, and drug sensitivity.

## Results

### Systematic comparison of Histone PTM and metabolite profiles

We first evaluated the correlation profiles between intracellular metabolite levels and histone PTM levels across 623 CCLE cell lines^36,37^ (**Figure 1A**; **S. Table 1**). We hypothesized that variation in levels of metabolites will lead to a corresponding change in levels of histone PTMs they potentially modulate. Among the top 20 metabolites with the highest absolute correlation value, we found several metabolites directly or indirectly linked with histone PTM levels, including AMP, pantothenate, NAD, and 1-methylnicotinamide (1-MNAM) (**Figure 1C**). Most of the top hits were related to the one-carbon metabolism pathway that produces nucleotides, glutathione, polyamines, and SAM^38^.

**Figure 1.**
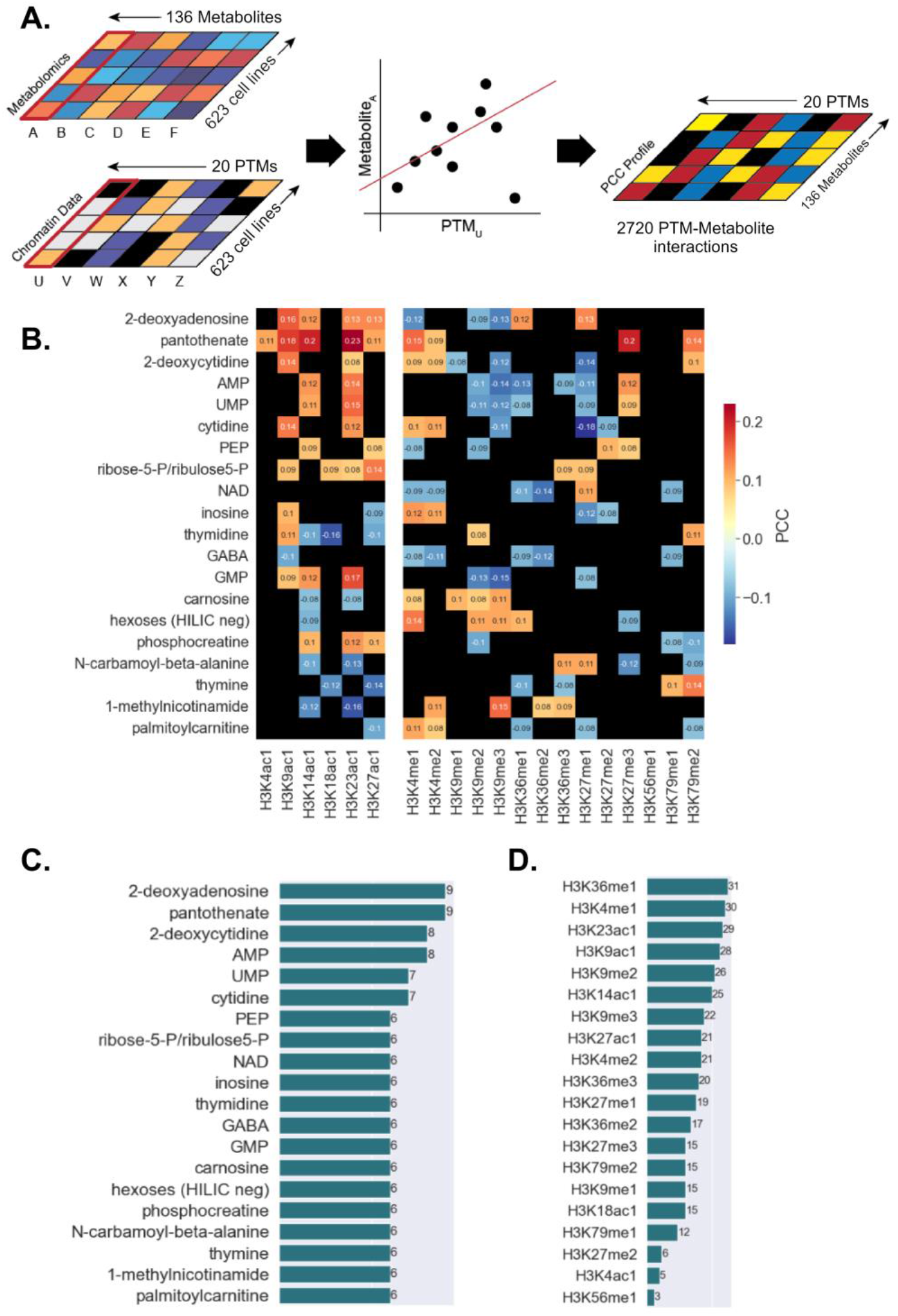
Metabolite and histone PTM correlation profiles across 623 cancer cell lines. **A)** Schematic of generating the Pearson’s correlation coefficient profile between metabolites and histone PTM data. Correlations are calculated between a PTM (example: PTM U) and metabolite (example: metabolite A) across all cell lines. This process is iterated across all metabolite and PTM combinations to generate a heatmap. **B)** Histone PTM and metabolite correlation profile for the top 20 metabolites with the highest number of statistically significant interactions (p-value < 0.05; black = non-significant). PTMs H3K9ac1, H3K9me1, H3K9me2, H3K9me3, H3K79me1 and H3K79me2 have known gene repressive actions. All other PTMs are known for gene activation. **C)** Number of statistically significant metabolites associated with a given histone PTMs. **D)** Number of statistically significant histone PTMs associated with a given metabolite. The top 20 metabolites were visualized.

1-MNAM is produced by Nicotinamide N-methyltransferase (NNMT) using SAM as the methyl donor and nicotinamide (NAM) as the substrate as a salvage mechanism^39^. We found that 1-MNAM is negatively correlated with histone acetylation while positively correlated with histone di- and trimethylation levels, specifically H3K4me2, H3K9me3, H3K36me2 and H3K36me3 (**Figure 1B**). Previous findings report that 1-MNAM upregulates histone deacetylase activity^40^, supporting our observation that 1-MNAM levels correlate with lower histone acetylation levels. The positive association between histone methylation and 1-MNAM levels was expected. However, it is interesting that this relationship is statistically significant only for selected lysine sites and is not associated with all methylation marks.

The levels of several nucleotides (NTs) were found to be positively correlated with histone acetylation. The correlation of these NT metabolites with histone PTMs is likely due to their connection with cellular growth. Given that global histone acetylation is generally associated with growth^41^, it is not surprising that these building blocks positively affect acetylation status. Ribose-5-P/ribulose-5-P from the pentose phosphate pathway was also positively associated with histone acetylation and is a necessary substrate for purine biosynthesis^42^.

Although the primary focus of this study is on metabolite-epigenetic interactions, we also found several significant associations between lipids and histone PTMs (**S. Figure 1**). To explore the impact that metabolites have on gene regulation, we grouped significant correlations by PTMs with known activating/repressing functions^43,44^ (**Figure 1B**). We did not find any metabolites with associations specific to activation or repressive marks alone.

We next sought to identify histone PTMs that are highly associated with metabolism. Our analysis revealed that H3K36me1, H3K4me1, H3K23ac1, H3K9ac1, and H3K9me2 were the top 5 histone PTMs that had the highest number of statistically significant associations (p-value < 0.05, FDR < 0.25) with metabolite levels (**Figure 1D**). Previous literature supports some of these top histone PTMs. For instance, several groups have identified the role of H3K9 and H3K4 methylation and acetylation on glycolytic activity^17,45^.

We performed extensive literature mining to confirm that we found biologically relevant metabolite/histone PTM interactions. We found 59 studies supporting several statistically significant interactions and their impact on histone PTM levels (**S. Table 2**). Many of these studies focused on metabolites that directly contribute to histone acetylation and methylation, including acetate, L-ascorbate, glucose, branched-chain amino acids, and fatty acids. Our correlation analysis also identified known epigenetic modulators and several novel metabolites with unannotated functions in epigenetic regulation (**Figure 1B**). Nevertheless, several factors may confound our correlation analysis, including the effect of tissue lineage and noise in omics measurements. We next correlated metabolite levels with epigenetic drug sensitivity data; we hypothesized that this orthogonal analysis could cross-validate metabolite-histone PTM interactions and uncover additional interactions.

### Systematic comparison of metabolite levels and epigenetic drug sensitivity

We analyzed data for 25 HDAC-, 5 HMT-, and 3 Histone demethylase (HDM)-inhibitors across 623 CCLE cell lines with drug sensitivity AUC values and metabolomics data^36,46^ (**Figure 2A**; **S. Table 3)**. In our analysis of the top 20 histone PTM drugs ranked by the number of significant metabolite correlations, we found 13 histone deacetylase (HDAC) and four histone methyltransferase (HMT) inhibitors (**Figure 2B**). Interestingly, we observed that the direction of the correlation (positive/negative) for each metabolite was the same between different drug classes (HDAC / HMT / HDM inhibitors). The exceptions to this observation were Repligen 136 and BRD-K80183349, suggesting that these drugs have different metabolic dependencies compared to other drugs within the same drug class.

**Figure 2.**
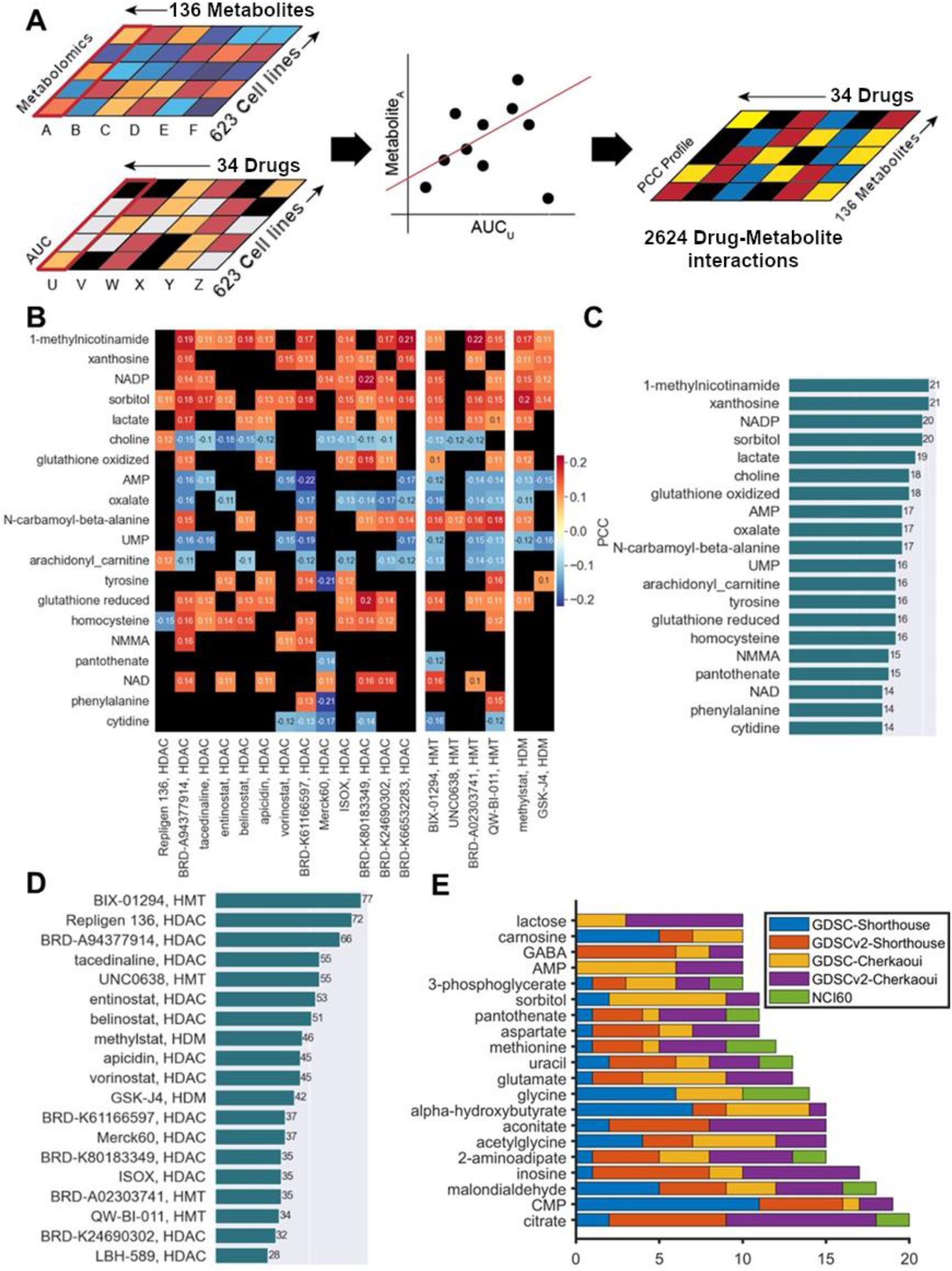
Metabolite and drug sensitivity correlation profiles across 623 cancer cell lines. **A)** Schematic of generating the Pearson’s correlation coefficient profile between metabolites and histone modifying drug AUC data. Correlations are calculated between a drug (example: drug U) and metabolite (example: metabolite A) across 623 cell lines. This process is iterated across all metabolite and drug combinations to generate a heatmap. **B)** Drug and metabolite correlation profile for the top 20 metabolites with the highest number of statistically significant interactions (p-value < 0.05, FDR < 0.25; black = non-significant). The x-axis is divided into three classes of inhibitors: HDAC, HMT, and HDM. **C)** Top 20 statistically significant metabolites (rows) associated with drugs. The bar chart illustrates the count of drugs significantly linked to a metabolite. **D)** Number of statistically significant histone-modifying drugs (x-axis) associated with a given metabolite. The top 20 drugs were visualized. **E)** Top epigenetic metabolites based on significant correlation with epigenetic drugs across multiple studies. Metabolomic data from Cherkaoui and Shorthouse *et al.,* studies^53,54^ were used to perform correlation analysis with epigenetic drug response data from GDSC and GDSCv2. The top 20 metabolites were visualized and the number of interactions with epigenetic drugs is shown in the x axis.

Overall, most drugs were significantly correlated with metabolites (**Figure 2C**). Both 1-MNAM and pantothenate appeared in the top 20 metabolites with the most interactions in the metabolites/histone PTM and metabolite/drug sensitivity analyses, suggesting that there is a significant epigenetic role for these two metabolites. Additionally, glutathione and homocysteine appear in the top 20 most connected metabolites, supporting the importance of redox homeostasis and one-carbon metabolism in modulating drug sensitivity. Most drugs that have statistically significant interactions with metabolites were HDAC inhibitors (HDACi), which may be partially attributed to the distribution of drugs available in the drug response dataset (**Figure 2D**).

We next analyzed synergistic and antagonistic relationships between drug/metabolite pairs. In this analysis, a positive correlation between metabolite level and drug AUC value implies that the metabolite/drug pair is in a potentially antagonistic relationship and vice-versa for a negative correlation (a high drug AUC value indicates greater cell viability). Our analysis revealed that 1-MNAM and homocysteine were associated with antagonism, while pantothenate was associated with synergism across all drug classes (**Figure 2B**). We also observed an antagonistic association between glutathione (GSH) with various epigenetic drugs. Increased levels of GSH have been associated with drug resistance in tumor cells through several mechanisms that support the redox environment of a tumor or that directly affect the drug^47^. GSH has also been shown to inhibit S-adenosyl methionine synthetase activity^48^, affecting DNA and histone methylation. Thus, the antagonistic association between GSH and histone PTM-modifying drugs could be attributed either to a direct influence on drug activity by modulating the redox balance or by limiting substrate availability. Finally, when considering novel and unexpected metabolite/drug associations, we found that sorbitol had an antagonistic association across all histone PTM drug types. Sorbitol is both an osmolyte and an activator of the c-Jun N-terminal kinase (JNK) pathway, which controls various processes, including proliferation and apoptosis after DNA damage^49^. Thus, sorbitol may modulate epigenetic drug potency in multiple ways, although the exact mechanism is not clear.

### Comparison of histone PTM levels and epigenetic drug sensitivity

To complement our metabolite-histone-PTM and drug sensitivity correlation analysis and ensure that we can capture biologically relevant associations with our dataset, we evaluated the correlation between histone PTM levels and histone PTM drug sensitivity (**S. Figure 2**). A negative correlation of PTM levels with drug sensitivity implies increased PTM levels are associated with decreased cell viability. We expected a negative correlation between histone acetylation levels and HDAC inhibitor AUCs, since HDAC inhibitors promote cell death via hyperacetylation. We found that our analysis supports our expectation and all statistically significant correlations were negative (**S. Figure 2D**; p-value < 0.05). We found that HDAC inhibitors - Vorinostat, Tacedinaline, and Apicidin, were the top three drugs with the highest association with histone PTM levels (**S. Figure 2A**).

In contrast to acetylation, the association between histone methylation levels and methylation-related drug sensitivity data is unclear (**S. Figure 2B**). We found several statistically significant positive and negative correlations between histone methylation levels and their respective HMT/HDM drug classes (**S. Figure 2C**), which was unexpected. HMT and HDM inhibitors had fewer statistically significant associations overall (**S. Figure 2A**). Finally, we examined which histone PTMs had the highest number of statistically significant associations with drug sensitivity (**S. Figure 2A & B; S. Table 4**). Among the acetylation PTMs, the most predictive lysine sites were H3K23, H3K14, H3K18, and H3K27.

### Consistency of metabolite and epigenetic drug sensitivity associations in other datasets

To assess the robustness and generalizability of these associations, we performed a similar correlation analysis using other large-scale cell line metabolomic profiling and drug sensitivity datasets from the literature. We first used data from the NCI60 cancer cell line panel, which has been extensively characterized using metabolomics and screened using a large panel of drugs^50–52^. Metabolite levels across the cell lines were correlated with the response of the same cell lines to epigenetic drugs from the CellMiner database. Associations from this NCI60 data overlapped significantly with those obtained from CCLE, higher than expected from random chance (hypergeometric p-value = 0.02) (**S. Table 3B**). We next used metabolomics data that profiled 180 different cancer cell lines in fixed growth conditions^53,54^. The metabolomics data was then correlated with epigenetic drug response data for the same cell lines from the genomics of drug sensitivity in cancer (GDSC) project^55,56^. This analysis also revealed a more significant overlap than expected out of chance with the associations obtained using CCLE data (hypergeometric p value = 0.0002) (**S. Table 3B**). Notably, several metabolites including pantothenate, sorbitol, methionine, and citrate, showed associations with epigenetic inhibitors across all these datasets (**Figure 2E**).

Overall, the correlation profiles revealed several important metabolite-PTM, metabolite-drug, and PTM-drug interactions across hundreds of cancer cell lines of different tissue lineages and genetic profiles. The findings from our correlation analysis were used to inform our drug profiling experiments.

### Inferring PTM, metabolite, and drug sensitivity interactions using statistical learning

While correlation analysis has identified important metaboloepigenetic relationships, correlation does not identify non-linear and combinatorial associations between many features. To address these issues, we leveraged three statistical learning methods that are simple to interpret to map the associations between metabolite levels, drug sensitivity, and histone PTM levels (**Figure 3**). We used the k-Top Scoring Pair algorithm (k-TSP), which measures ratios of metabolites. We used this approach since many cellular processes are sensitive to relative levels of metabolites (e.g., NADH/NAD) rather than absolute levels. In addition, LASSO and stepwise regression were used to get the minimal number of metabolites in combination that best predict either a given histone PTM or drug AUC value (**Figure 3A**).

**Figure 3.**
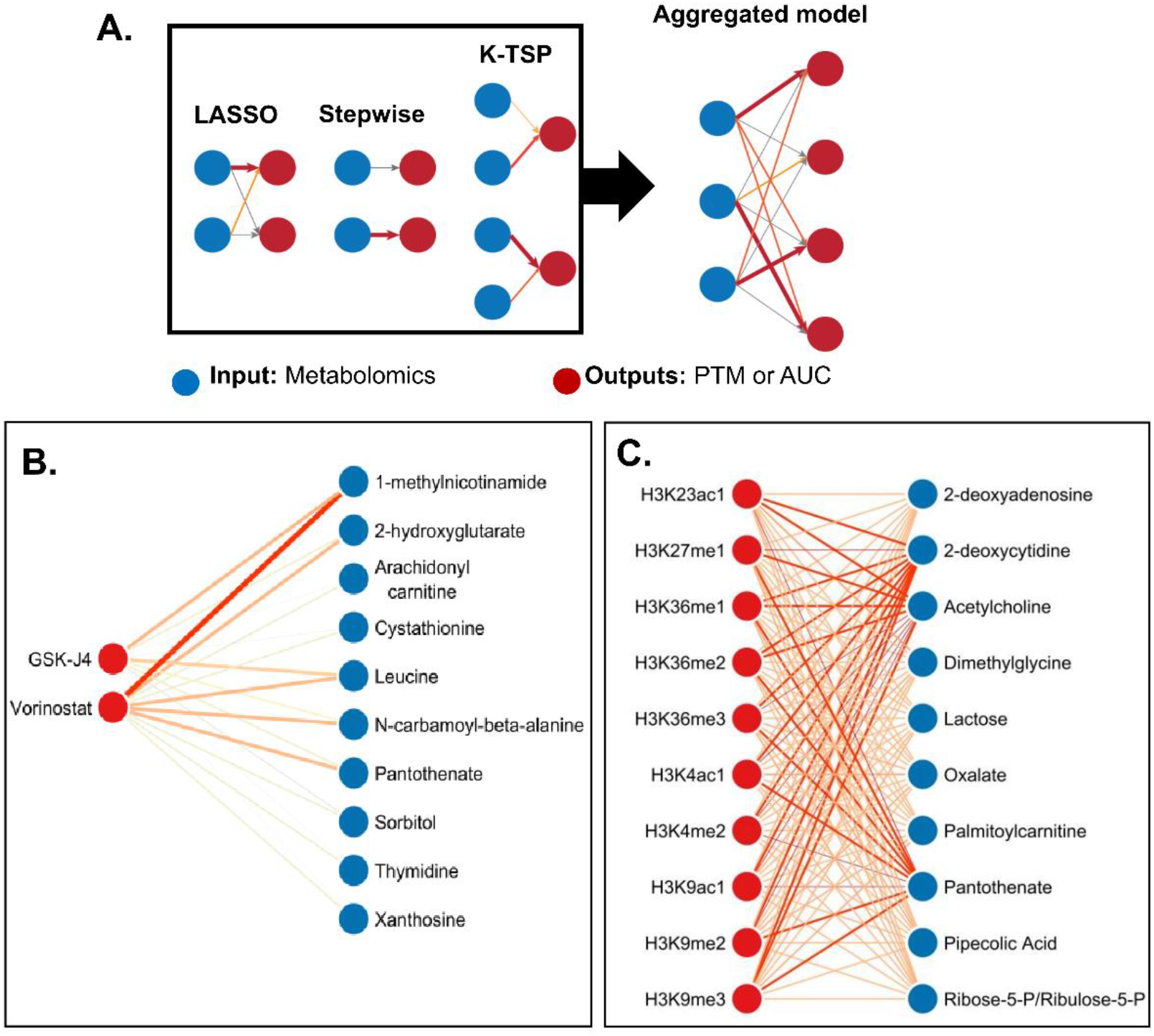
Inferring metabolite-PTM-drug sensitivity interactions using machine learning. **A)** Model schematic. Three machine learning models were trained on metabolomics data (input: metabolomics, blue nodes) to predict histone PTM levels and drug AUC values (output; red nodes). The absolute value of the coefficients for the LASSO and stepwise regression models and the Matthew’s correlation coefficient of the K-TSP true/predicted outputs were combined over the union using a product operation. The ensemble-model coefficients were weighted by the number of experiments that intersected across the models. The edge color (gray: weak to red: strong) indicates the strength of the relationship between the two nodes. Top interactions consistent across multiple methods are visualized in panels B and C. See **S. Tables 8 and 9** for a complete list of all interactions. **B)** Top ensemble-model relationships between metabolite levels and GSK-J4 and Vorinostat drug AUC levels. Red nodes indicate epigenetic drugs. Blue nodes indicate metabolites. **C)** Top ensemble-model relationships between metabolite levels and histone PTM predictions. Red nodes indicate PTMs. Blue nodes indicate metabolites.

We first analyzed individual machine-learning model findings as each model uncovered a distinct set of interactions. LASSO is a powerful approach that performs feature selection and regression simultaneously. When working with large datasets, LASSO can identify a small subset of variables that are important and predictive. We applied LASSO feature selection to identify top metaboloepigenetic interactions (**Methods**). We used the LASSO coefficients to construct a bidirectional directed network to quantify the interactions between histone PTMs and metabolite levels (**S. Figure 3; S. Table 5**).

We found several metabolites that are known to support both acetylation and methylation processes. Methionine, threonine, and 1-MNAM from one-carbon metabolism were associated with the H3K4me1, while phosphorylated sugars from glycolysis (F1P/F6P/G1P/G6P) and branched-chain amino acids were linked with H3K14ac1 (**S. Figure 3**). These are consistent with prior studies linking these metabolites with histone PTMs^57,58^. Other notable metabolite/PTM interactions include the positive associations between pantothenate and H3K4me1, and the negative association between kynurenic acid and H3K23ac1. Notably, these interactions were not identified in our correlation analysis. This is because correlation fails to account for combinatorial interactions, while LASSO can account for multiple factors impacting a variable of interest, in this case, multiple metabolites interacting with a histone PTM. These suggest that the histone PTMs are potentially sensitive to relative levels of these metabolites in relationship with other metabolites.

While the LASSO interaction network is intuitive, the large number of associations is still a barrier to interpretation. Therefore, we trained simpler stepwise-regression models that only considered a single metabolite, two metabolite, or three metabolite models that best-predicted histone levels (**S. Figure 5; S. Table 7**). Across all our models, we found that pantothenate, leucine, and 1-MNAM were top predictors of histone PTM levels and drug sensitivity.

Since correlations, LASSO, and stepwise regression models assume a univariate linear contribution to each output, we used the k-TSP model to assess bi-variate interactions. This modeling approach evaluates the interactions two metabolites have on a given histone PTM with a given probability. We also assessed predictions from other machine learning models and found k-TSP to provide both relatively high accuracy and interpretability (**Methods**). k-TSP identified that the relative levels of 1-MNAM with lactose and kynurenic acid contributed strongly to histone PTM predictions, specifically H3K23ac1 and H3K9me3. Further, our analysis revealed that 1-MNAM together with palmitoylcarnitine, and S-adenosylhomocysteine with aconitate, are predictive of H3K9me3 levels (**S. Figure 6D-E**). We also evaluated which PTM pairs are predictive of metabolite levels. We found that H3K23ac1/H3K9me3 ratio and H3K14ac/H3K79me2 ratio were predictive of 1-MNAM (**S. Figure 6A-C**).

We combined the coefficients from all these models into a single ensemble model to identify the most robust interactions under different statistical assumptions. 2-deoxycytidine, acetylcholine, and pantothenate had the highest associations with several histone PTM levels in the ensemble metabolite-PTM model (**Figure 3B; S. Table 8**). Pantothenate, which is required for coenzyme A biosynthesis^59^, was highly predictive of H3K4ac and, to some extent, H3K9ac and H3K23ac across multiple models. Surprisingly, a strong relationship emerged between pantothenate and several histone methylation marks suggesting that it may have a cross-functional regulatory role with histone methylation. However, there was no reported experimental evidence supporting this finding.

We next looked at two epigenetic drugs that were highly correlated with metabolite levels. These two drugs were Vorinostat, an HDAC inhibitor that targets most HDACs, and GSK-J4, an H3K27me2/3 demethylase inhibitor^60,61^. Vorinostat is used for treatment for cutaneous T-cell lymphoma and is in clinical trials for several cancers, while GSK-J4 is being explored as a treatment for gliomas^62^. Notably, we observed a strong association between Vorinostat and 1-MNAM (**Figure 3C; S. Table 9**). A previous study has shown that 1-MNAM stabilizes the deacetylase enzyme SIRT1^63^ and may also impact the activity of other deacetylases. Our machine learning approach supports this hypothesis by predicting its association with Vorinostat.

GSK-J4 was also associated with 1-MNAM and leucine across multiple machine learning models (**Figure 3C; S. Table 9**). Of note, we found a synergistic link between GSK-J4 and kynurenic acid (**S. Figure 4; S. Table 6**). Kynurenic acid is known for its anti-inflammatory and immunosuppressive functions that support cancer progression^64^. Kynurenic acid has been shown to impact histone methylation patterns in microglial cells to support anti-inflammatory phenotypes^65^. However, the role of kynurenic acid in modulating epigenetic drug sensitivity and cancer epigenome has not been investigated.

We summarize our findings for metabolite/PTM interactions (**S. Table 10**) and metabolite/drug interactions (**S. Table 11**) that are common across multiple machine learning and statistical models. While our analysis revealed that metabolites 1-MNAM and kynurenic acid were highly associated with different histone methylation and acetylation marks, Li *et al.,*, 2019 found that strong lineage effects are associated with these metabolites^36^. To confirm whether this is a real biological signal, we next experimentally assessed the impact 1-MNAM and kynurenic acid had on epigenetic drug sensitivity.

### Validating metabolite and epigenetic drug interactions

Our model results identified 1-MNAM, pantothenate, kynurenic acid, glutathione, and sorbitol as important predictors of both histone PTM levels and epigenetic drug sensitivity. To validate our predictions, we performed a metabolite + drug combination screen to assess how these metabolite-drug interactions impact cell proliferation (**Figure 4A**).

**Figure 4.**
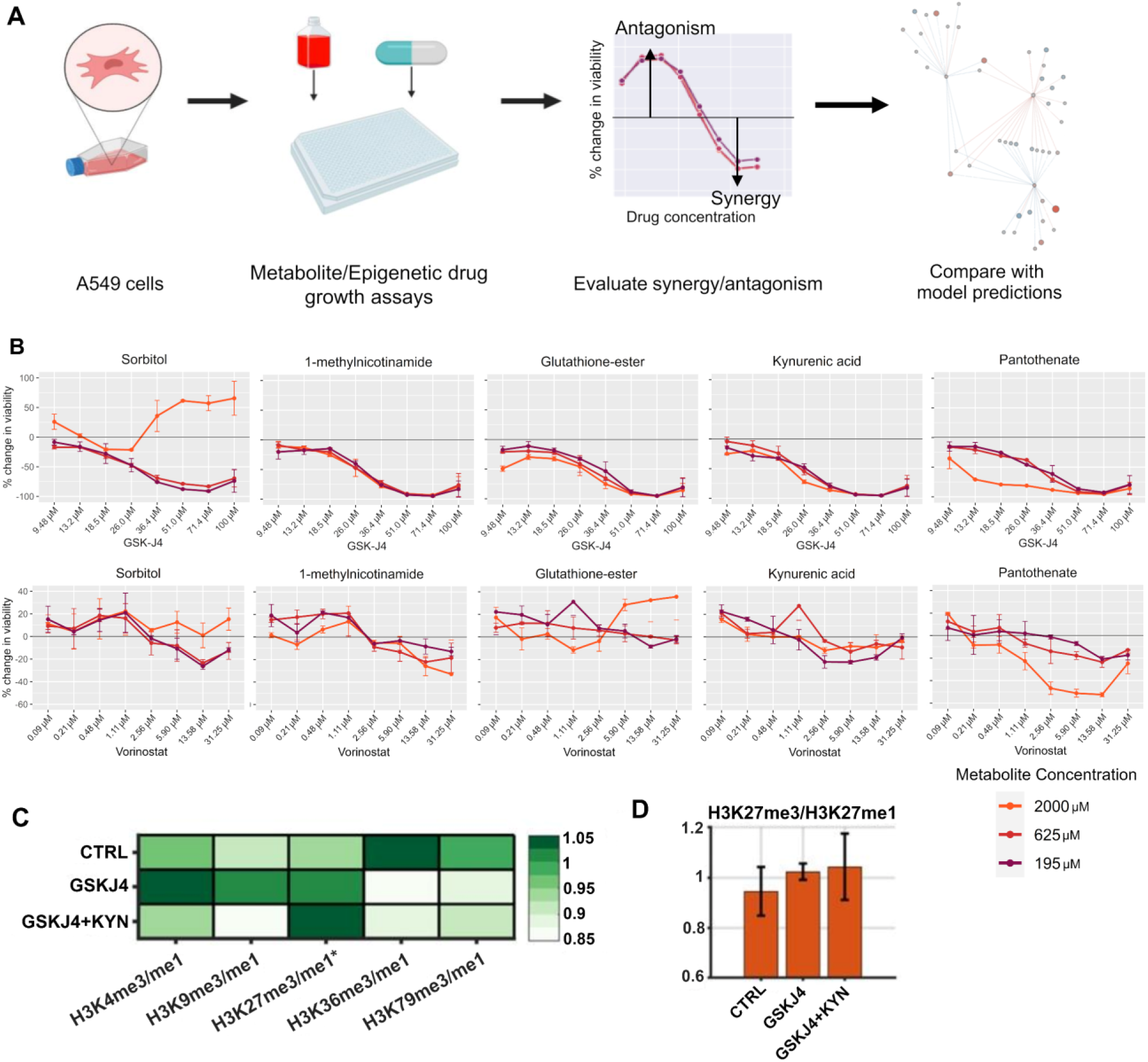
Metabolite/drug dose-response curves to validate model predictions for Vorinostat and GSK-J4 in A549 cells. **A.** Schematic of the experimental procedure. **B.** Percent viability curves (treatment/control) for five metabolites + epigenetic drugs Vorinostat and GSK-J4. Control was the drug-only condition. The black line indicates no difference between treatment and control. Data points were averaged across replicates (N=2 treatment replicates; N=12 control replicates). Negative percent values below the black bar indicate synergy, while positive percent values indicate antagonism. **C** and **D.** Relative change in five major H3 methylation marks (trimethylation/monomethylation) for control (no treatment), GSK-J4 treatment and the combination of GSK-J4 and Kynurenic acid treatment. Only H3K27 shows a statistically significant increase in levels for treatment compared to control. The H3K27 data is highlighted in the bar chart (average of two replicates; t-test p-value = 0.056 for control vs combination treatment)

We ran our metabolite/epigenetic drug screen using A549 as the cell line model with epigenetic drugs Vorinostat and GSK-J4. We ran the synergy/antagonism screen with an eight-point drug concentration profile with three concentrations of metabolite supplementation (**Methods**). When we evaluated the dose-response curves using DMSO as the control, we found a dose-dependent response across all metabolite and drug-treated samples (**S. Figure 8**).

Notably, GSK-J4 demonstrated significant synergy across all metabolites except sorbitol, which exhibited an antagonistic effect at high concentrations and synergy at a lower dose (**Figure 4B**). When we evaluated Vorinostat/metabolite combinations, we found a neutral effect with 1-MNAM, a synergistic effect with pantothenate, and mild antagonism with both sorbitol and glutathione. The synergistic effect pantothenate has on both the HDAC inhibitor and the histone demethylase inhibitor suggests that it has multiple roles in epigenetic regulation and cellular survival that have not been reported in the literature. Reduction in intracellular glutathione levels is known to increase sensitivity to Vorinostat^66^, supporting our antagonistic interaction of glutathione with Vorinostat.

Further, we found that high sorbitol levels antagonized the efficacy of both drugs. While our experimental results agree with our model predictions, we must acknowledge that our analyses have confounding factors. The metabolomics data we used in our correlation analysis measured internal levels of sorbitol, while our assay supplements sorbitol to the medium. External sorbitol application in cell culture results in mild osmotic stress^67^, which may affect cell proliferation.

To tease out dose-specific interactions, we assessed significant pairwise metabolite and drug interactions and their impact on cell viability (**S. Figure 11**). To filter out noise from the dataset, we applied a p-value threshold to identify interactions with a discernible difference in the average cell count between treatment and control (two-sample T-test p-value threshold < 0.05). Our p-value filter found significant antagonistic interactions between Vorinostat with both 1-MNAM and glutathione at low metabolite doses (**S. Figure 11B**), which was also predicted from our correlation analysis (**Figure 2B, 3C**).

Most metabolite combinations showed a statistically significant synergistic effect with GSK-J4 (**S. Figure 11A**). On average, interactions were significant at the higher drug doses and across all three metabolite concentrations for all metabolites. Pantothenate and GSK-J4 show a dose-dependent synergistic relationship on cell viability, especially at higher levels of pantothenate. Further, to assess the effect of the cell line model used on these interactions, we confirmed the strong synergistic interaction between kynurenic acid and GSK-J4 in HCT116 cells, which are derived from a different tissue lineage (colon) compared to A549 cells (lung) (**S. Figure 10**). Of note, we had used a higher concentration for metabolites (mM range) compared to drugs (µM range) since that provided a clear phenotype without impacting cell growth, consistent with prior studies on drug-nutrient interactions^32,68,69^. We performed an additional experiment at lower levels of metabolites to check if the synergistic interactions occur at lower doses (**S. Figure 9**). We used the bliss statistical metric that controls for both drug and metabolite effects at each dose to quantify the interactions (**Methods**) and this metric predicts consistent synergy between Kynurenic acid and pantothenate with both GSK-J4 and vorinostat. The metabolites on their own do not have an inhibitory effect in the range tested. Hence, the synergy is directional, the metabolites enhance the potency of the drugs and not vice versa. Therefore, the percentage change in drug potency upon metabolite addition is reported throughout this study as it is more straightforward to interpret.

The statistical learning models allowed us to identify significant interactions that were missed by correlation analysis, notably for 1-MNAM, glutathione, kynurenic acid, and pantothenate with GSK-J4. In addition to sorbitol/GSK-J4, sorbitol/Vorinostat, and glutathione/Vorinostat, we predicted kynurenic acid having a synergistic interaction due to its negative LASSO coefficient with GSK-J4 (**S. Figure 4**). Our analyses did not show any significant association between kynurenic acid and Vorinostat, which was also captured in our assay results at high doses. Interestingly, our correlation analysis suggested that 1-MNAM was antagonistic to

Vorinostat. However, the relationship is hard to determine as the combination shows antagonism at lower drug doses and synergy at higher doses (**Figure 4B**). We summarize experimentally determined synergy/antagonism between metabolite and drug in **Table 1**.

**Table 1.** Summary of experimentally observed synergy/antagonism between drug metabolite pairs. (predictions from correlation and ensemble model that match with experimental data are bolded) (Neutral – no interaction or inconclusive).

| Metabolite | Drug | Experimentally determined synergy/antagonism |
| --- | --- | --- |
| Sorbitol | GSK-J4 | **Antagonism** |
| 1-MNAM |  | **Neutral** |
| Glutathione |  | Synergy |
| Kynurenic Acid |  | **Synergy** |
| Pantothenate |  | Synergy |
| Sorbitol | Vorinostat | **Antagonism** |
| 1-MNAM |  | **Neutral** |
| Glutathione |  | **Antagonism** |
| Kynurenic Acid |  | **Neutral** |
| Pantothenate |  | Synergy |

### Dissecting mechanisms underlying synergy between kynurenic acid and GSK-J4

The interaction between kynurenic acid and GSK-J4 is especially surprising as there are no reported interactions between kynurenic acid with H3K27me3 or GSK-J4. GSK-J4 affects H3K27 methylation by inhibiting the demethylase KDM6A and KDM6B^61^. To evaluate if kynurenic acid also synergizes with GSK-J4 epigenetically, we measured the epigenetic effects of kynurenic acid and GSK-J4 by experimentally measuring different histone H3 modifications in A549 cells. As expected, treatment with GSK-J4 increased H3K27me3 levels (**Figure 4C**). Notably, treatment with both kynurenine and GSK-J4 further increased H3K27me3 (p-value = 0.054) and no other H3 modification (p-value > 0.05) (**Figure 4D**). Thus, their synergistic growth inhibition may be mediated through synergistic hypermethylation at H3K27me3.

GSK-J4 is being explored for treatment of gliomas^70^. Further kynurenic acid can cross the blood brain barrier and has been associated with glioma pathophysiology^71^. Hence its interaction with GSK-J4 might be clinically relevant for treatment of these brain cancers. To study the potential clinical relevance of this interaction, we analyzed Chip-seq data from GSK-J4 treated diffuse intrinsic pontine glioma (DIPG) cells^70^. This revealed that Indoleamine 2,3-Dioxygenase 1 (IDO1) is the most differentially methylated gene at H3K27 upon GSK-J4 treatment^70^ (log fold change = -1.3, p-value = 0.015 for treatment vs control). Since H3K27 trimethylation is a repressive mark, the corresponding IDO1 transcript is significantly upregulated after GSK-J4 treatment (log fold change = 0.41, p-value = 0.054 for treatment vs control). Kynurenic acid is derived from tryptophan metabolism and IDO1 represents the rate limiting step of its formation^72^. Metabolomics data from glioma cell lines that harbor a H3K27 mutation, which prevents methylation, also revealed that kynurenine and tryptophan are significantly downregulated due to the mutation compared to the wild type (log fold change = -0.22 and -0.24, p-value = 0.04 and 0.008 for kynurenine and tryptophan respectively, mutant vs wildtype)^73^. These results suggest that kynurenic acid metabolism is regulated by H3K27 trimethylation and perturbing this PTM through GSK-J4 increases kynurenine levels. Thus, the synergistic growth inhibition of kynurenic acid and GSK-J4 may also be mediated through H3K27 trimethylation leading to imbalances in kynurenine metabolism. Analysis of the PRISM drug screening database provides additional independent evidence for these interactions^74^ (**S. Figure 12**). Levels of kynurenine in brain cancer cell lines significantly correlate with the corresponding potency of GSK-J4 (p-value = 0.035). GSK-J4 potency across brain cancer cell lines also correlates with the expression and dependency of IDO1 enzyme (p-values = 0.08 and 0.002 respectively) (**S. Figure 12**). These results further support the predicted interaction between kynurenine metabolism, H3K27 trimethylation and GSK-J4.

### Tracing metabolite-epigenetic changes during the epithelial-to-mesenchymal transition (EMT)

As a proof of concept of how the inferred network can be used to understand metaboloepigenetic interactions in various contexts, we applied our model to the EMT system. EMT is a reversible developmental process that facilitates the transition from an epithelial cell to a mesenchymal cell with stem-cell-like properties^75^. Extensive metabolic and epigenetic changes occur during EMT, which may contribute to tumor heterogeneity, drug resistance, and metastasis^76^. How metabolic and epigenetic processes interact with each other to promote EMT is unclear.

We tracked changes observed in metabolomics and histone proteomics data in A549 cell line undergoing EMT^77,78^ (**Figure 5; S. Figure 7**). Overlaying the histone PTM data onto the metabolite/PTM network showed that most changes occurred during the first 24-hour period. H3K4me2 showed the largest decrease, and H3K9ac showed the largest increase in levels relative to the start of the experiment (**S. Figure 7**). We next applied our trained LASSO models to quantitatively predict changes in PTM levels over time using time-course metabolomics and histone PTM proteomics data. The data was centered on the same scale as the CCLE training set (**Methods**). We evaluated the quality of our predictions by comparing the mean-squared error (MSE) between the predicted values against the true PTM levels (**Figure 5B**). Overall, most histone PTMs were predicted to have a low error, showing that our LASSO model was generalizable across datasets. The two histone PTMs our LASSO model could not predict well were H3K4me2 and H3K27me1, suggesting heterogeneity and context-specificity of their expression. To understand the metabolites driving these predictions, we next visualized the metabolomics data with the LASSO metabolism-PTM network (**Figure 5C**). The most pronounced changes in epigenome-predictive metabolite levels were increased 1-MNAM, glycolytic metabolites F1P/F6P/G1P/G6P and dimethylglycine (DMG). These observations corroborate previous studies, where increased glycolysis and one-carbon metabolism were associated with EMT^79–81^. To complement the metabolite-PTM analysis, we next focused on the metabolite-drug interaction network. This revealed that the epigenetic metabolites 1-MNAM and 2-hydroxyglutarate increased from 24-72 hours based on the metabolomics data, which are associated with Vorinostat drug sensitivity (**Figure 5A**). Vorinostat displays a dual response of inducing and inhibiting EMT in a context-dependent manner^82,83^. Our model captures the context-specific heterogeneity in Vorinostat response based on metabolic state, especially with 1-MNAM, which shows dose-dependent interaction with Vorinostat.

**Figure 5.**
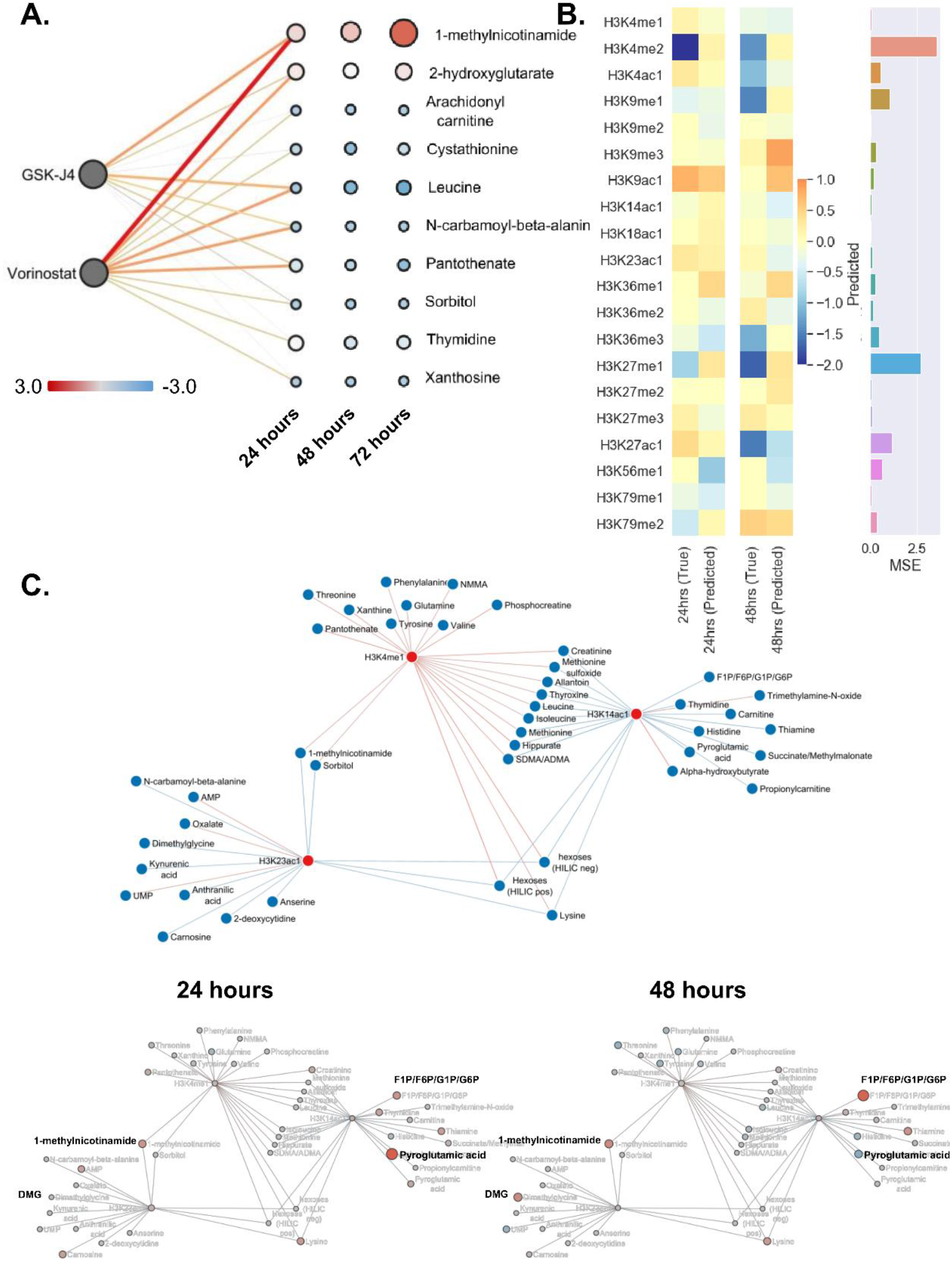
Assessing metabolic and epigenetic interactions during the Epithelial-to-Mesenchymal (EMT) transition. **A.** EMT time-course metabolomics^77^ was measured from 0-72 hours, and the relative values are overlaid onto the ensemble-model network from Figure 3. Log2 fold change (LFC) of the metabolomics (24 hours / 0 hour) is shown, where red is a positive LFC, gray is an LFC close to 0, and blue is a negative LFC. The size of the metabolite node is the absolute value of the LFC. Metabolites that did not intersect between the CCLE and the metabolomics data include arachidonyl carnitine, xanthosine, sorbitol, and N-carbamoyl-beta-alanine. **B.** LASSO predictions on Nakasuka *et al.,*, 2021 metabolomics data to predict^78^ histone levels during EMT. Barplot shows mean-squared error (MSE) for both 24 and 48hrs. Heatmaps show predicted values and LFC for histone PTM data for 24 and 48 hrs. **C.** LASSO network showing the relationship between metabolomics and histone PTM LFC. Key metabolites are highlighted.

## Discussion

Here we use a multi-omic statistical learning framework to study how cellular metabolism impacts histone PTMs and epigenetic drug sensitivity. Nutrients and metabolic state can significantly impact drug sensitivity in cancer and other diseases. For instance, dietary methionine restriction increased sensitivity to radiation and chemotherapy in mouse models^84^. However, the interactions between metabolites and epigenetic drugs have not been systematically characterized.

Our study leverages LASSO, stepwise, and k-TSP algorithms to capture the non-linear and combinatorial interactions between metabolites and histone PTMs. While analysis of drug sensitivity data alone captured some synergistic/antagonistic interactions, augmenting with histone PTM levels helped us comprehensively identify additional epigenetic modulators. One example was pantothenate, which showed strong correlations and associations with histone PTM levels but not with the drug AUC data. When tested experimentally, we found strong synergy with both GSK-J4 and Vorinostat. In addition, choosing data mining methods with different statistical assumptions like LASSO and kTSP allowed us to comprehensively capture interactions that each technique would have missed individually. Our data suggests that PTMs are potentially sensitive to the relative levels of metabolites to other metabolites. This diverse range of metabolic sensitivities for various histone PTMs may be related to the number of enzymes regulating each PTM, their kinetic properties, and their interactions with metabolic enzymes.

Our data suggests that histone PTM levels can be influenced by several metabolites that are not known to be direct substrates for histone writers and erasers. Through a literature search, we identified 27 metabolites across 59 independent studies that had prior known effects on histone acetylation and methylation levels (**S. Table 2**). Our model predicted novel metabolite-PTM and metabolite-drug interactions, such as kynurenic acid with GSK-J4. Additionally, our ensemble-model found that 2-hydroxyglutarate (2HG), an oncometabolite produced from IDH1/2mt^85,86^, was associated with Vorinostat sensitivity. A recent study reports that IDH1-mutant cancers are sensitive to Vorinostat by suppressing homology-directed DNA repair in cells producing 2HG^87^. Thus, these independent studies support our model’s predictions.

Kynurenic acid was predicted to have a synergistic relationship with GSK-J4, which we confirmed experimentally (**Figure 4 & S. Figure 4**). Further, experimental measurement of H3 methylation levels suggest that their synergistic growth inhibition may be mediated through synergistic hypermethylation at H3K27me3. While the observed changes in k27me3 are statistically significant, the effect is rather small (10% change). Even the study that originally reported GSKJ4 shows similar magnitude of change in total K27me3 levels (5-10%), likely because GSKJ4 is a highly specific inhibitor^88^. Hence our experiments reflect the expected change in k27me3 levels. The inhibitory effect is likely due to an increase in methylation mark at specific loci regulated by JMJD3/UTX, the enzymatic target of GSKJ4, which affects H3k27Me3/Me1 ratio. Analysis of public chip-seq datasets after GSK-J4 treatment revealed that kynurenic acid metabolism is regulated by H3K27 methylation, further supporting the mechanistic link between kynurenic acid, GSKJ4 & H3K27 methylation. These results overall suggest a potential feedback regulation between metabolites and the corresponding PTMs they influence. The synergistic interaction with GSK-J4 may have clinical implications for treating gliomas that frequently exhibit mutations in H3K27 and dysregulation of kynurenic acid pathway^71,89,90^.

Pantothenate, a precursor of CoA biosynthesis^59^, showed a strong correlation with PTM levels and a synergistic effect in combination with GSK-J4 and Vorinostat. Manipulating acetyl-CoA levels can influence the degree of protein acetylation, which supports our experimental results, with pantothenate having a synergistic effect with Vorinostat. However, its synergy with GSK-J4 needs further experimental evaluation and may be mediated via EZH2, a H3K27 methyltransferase^91^. Acetylation of EZH2 requires acetyl-CoA thus potentially linking this metabolite with H3K27 methylation. Consistent with this, the surprisingly positive correlation between pantothenate and H3K27me3 in our dataset is lost in EZH2 loss of function mutant CCLE cell lines (R = -0.44, N = 16 cell lines), and is maintained in cell lines with gain of function mutants (R = 0.51, N = 9 cell lines).

Our model also found that 1-MNAM interacts with various histone PTMs, which may explain its diverse biological effects. 1-MNAM enhances the mesenchymal phenotype in cancer cells^92,93^ by increasing the deacetylase enzyme SIRT1^63^. A recent study found that 1-MNAM is also a critical immunomodulator within the tumor microenvironment^94^ and infectious diseases such as COVID-19^95^.

A limitation of our study is that we did not probe the exact mechanism underpinning these inferred metabolite/PTM/drug interactions. However, we do find that these interactions are robust across hundreds of cell lines, and their mechanistic connections can be reasonably explained. Our approach prioritized the strongest metabolite/PTM and metabolite/epigenetic drug interactions, which were further scrutinized using the consolidated result of multiple machine learning models with different statistical assumptions and model complexity. Finally, the top predictions that passed these analytical thresholds showed synergistic/antagonistic effects in our drug/metabolite experiments. While the experimental validation in two cell lines is still limited, we observe that the interactions were observed across hundreds of cell lines in CCLE and were seen in independent datasets from GDSC and NCI, and during EMT. Our ensemble model also correctly recalls several known interactions reported in literature. These suggest that the statistically inferred associations may represent real metaboloepigenetic interactions. Another limitation is that some of these interactions could be influenced by common oncogenic mutations. However, a previous analysis of metabolites and common genomic alterations like P53 mutations in the CCLE dataset found a very weak correlation between the two^36^. Consistent with this, we do not find a strong genetic association with the five metabolites and two drugs that were used for experimental validation (**S. Table 12, 13 and 14**).

Overall, our model identified critical metabolite-PTM and metabolite-drug interactions that were previously unidentified. These results provide new insights into metabolic control of the epigenome, and call attention to the impact of nutrient conditions on epigenomic assays, which is usually not considered. Our study can also shed light on the epigenetic effects of metabolites and other nutrients within the tumor microenvironment (TME). TME is a critical regulator of tumor progression and plays a role in acquiring drug resistance to various chemo- and radiation therapies^96–98^. The methodology outlined in this study can enable precision medicine applications. For instance, metabolomics can be used to generate hypotheses on epigenetic alterations and help prioritize epigenetic therapies. Furthermore, metabolite modulators of epigenetic therapies can be used to design specialized diets that enhance drug efficacy. HDAC inhibitors and epigenetic therapies are being explored for numerous cancers, immune- and neurological-diseases, and our results are potentially relevant to these diseases.

## Methods

### CCLE Data Preprocessing

Metabolomics^36^ and H3 global chromatin profiling^37^ data were obtained from the Cancer Cell Line Encyclopedia (CCLE), while drug sensitivity data^46^ was obtained from the Cancer Therapeutics Response Portal (CTRP). We used Broad Institute IDs to map across the various datasets, resulting in 623 cancer cell lines intersecting across all datasets. The data was transformed into a Z-score prior to downstream analyses for the metabolomics and global chromatin profile data. The CTRP data was left as-is for the correlation analysis or scaled using a minimum-maximum scaler function across each drug for machine learning to reduce the data dimensionality by its range. K-nearest neighbor imputation was performed to fill in missing values using the scikit-learn library and default arguments.

We chose to reduce the dimensionality of the datasets by focusing on the interpretability of our results. Histone PTMs with modifications at multiple lysine sites or no modifications were removed from the global chromatin profile dataset for most analyses except the K-TSP algorithm, reducing the number of PTM features from 42 to 20 histone PTMs. In the metabolomics dataset, lipids (triacylglycerides, ceramides, diacylglycerides, sphingomyelin, cholesterol esters, lysophosphatidylethanolamine, lysophosphatidylcholine, and phosphatidylcholine) were removed/deprioritized (**S. Figure 1**), resulting in 136 metabolites. For the CTRP drug area under the curve growth (AUC) data, we focused on drugs that affect histone PTMs, using the string ‘histone|sirtuin|HDAC|demthylase|methyltransferase’ as the filtering criteria to select drugs while filtering out drugs that affected DNA epigenetics. This reduced the number of drug features from 481 small molecules to 34 histone-modifying drugs.

### Estimating pairwise associations between histone PTMs, metabolite levels, and drug AUC values using machine learning

We initially trained nine machine learning models of varying model complexity to evaluate their ability to infer the relationship between histone PTM levels, metabolite levels and drug AUC values. Specifically, we evaluated ordinary least squares, logistic regression, ridge regression, LASSO, K-Top Scoring Pairs (KTSP), decision trees, random forest, gradient boosting, and support vector machines. Based on the trade-off between model accuracy, simplicity, and interpretability, we selected LASSO and KTSP as the regression and classification algorithms used for downstream analyses. In our LASSO models, we prioritized interactions within the network using a Pearson Correlation Coefficient p-value cutoff of 0.05 and chose the strongest interactions. Finally, we performed univariate analyses using 1-feature, 2-feature, and 3-feature stepwise regression models.

We included the K-Top Scoring Pairs (KTSP) method (Yoon & Kim, 2010), which is not commonly used in machine learning, but may be relevant in this study because of its ability to detect pairwise interactions. It inherently performs dimensionality reduction to select the top pairs of histone PTMs predictive of metabolite levels and vice-versa. A threshold of 0 was used to label the metabolite or histone PTM classified as upregulated (positive values) or downregulated (negative values). We performed additional feature engineering, including the sum and variance of histone PTM levels and the cell line growth doubling time to evaluate the impact of these features on metabolite or histone PTM levels. We trained k-TSP models on all histone PTM features and 136 non-lipid metabolites. An example of our implementation of the k-TSP algorithm is shown below:

For PTM pair P_A_, P_B_:

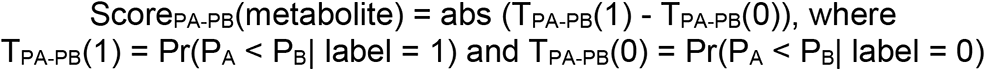

We performed a Bayesian hyperparameter optimization procedure for each model with 30 sets of hyperparameters to obtain the best model that maximized the Pearson Correlation Coefficient for regression-based tasks and the Matthews Correlation Coefficient (MCC) for classification-based tasks. Overall model metrics are reported as the average value from 10-fold cross-validation.

We used the Benjamin Hochberg approach to correct for multiple hypotheses and obtain a False Discovery Rate (FDR). We decided to use a liberal FDR threshold of 0.25 & p-value < 0.05 for all correlation analyses to obtain a significant number of interactions for comparison with other methods, which enabled us to aggregate the results of multiple machine learning models and infer the strongest associations. However, only the topmost correlations (p-value << 10^-3^) were highlighted for discussion.

### Combining multiple machine learning models into a single ensemble-model

To simplify model interpretation and identify the most important metabolite-drug and metabolite-PTM associations, we consolidated our LASSO, K-TSP, and stepwise regression models to get an ensemble-model. Specifically, we performed an outer join multiplication operation between the LASSO and stepwise regression coefficients. Because the K-TSP algorithm calculates unique sets of coefficients for each metabolite K-mer associated with a drug or PTM value, that limited our ability to incorporate coefficients into this ensemble-model. To resolve this issue, we took the Matthews correlation coefficient (MCC) between the predicted and true outputs as multiplicative weights with the LASSO/Stepwise regression product across all drug/PTM values. It is a common strategy in ensemble machine learning to weight each individual learning method based on its accuracy^99^. Here we use the Matthews correlation coefficient (MCC) as the accuracy metric and used that to weigh the coefficients of the kTSP model. The kTSP is a binary classifier for a set of paired entities (metabolites, histone PTMs). We found that the class distribution is imbalanced for k-TSP classes - more zeros than ones. Thus, the MCC is appropriate as the accuracy metric, as it is a fair way to take into consideration not only true positives and negatives, but also the false positives and negatives. MCC is well known to be a robust metric of accuracy and hence it was used as the multiplicative weight^100^. The final product, *θ* was used for all downstream analyses.

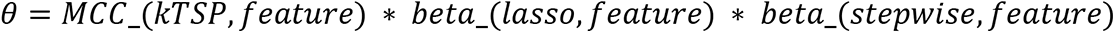

The resulting metabolite-drug and metabolite-PTM tables were still relatively high dimensional and comprised 136 metabolites. To simplify our results into a top 20 metabolite list, we took the sum of the consolidated coefficients across the drug, metabolite, and PTM dimension, which was used as our importance metric. Then, we took the top and bottom ten metabolites, resulting in a list of 20 metabolites for each association. We visualized two drugs of interest and the top/bottom 5 PTMs (for 10 PTMs).

### Drug/Metabolite/PTM interaction networks

We constructed bidirectional directed networks using the LASSO regression coefficients to study histone PTM / metabolite and metabolite/drug AUC relationships. The edge color represents the sign of the regression coefficient, while the edge size represents the magnitude of the LASSO coefficients, where values larger in magnitude indicate stronger relationships. Pairwise relationships with a value of zero indicate no association. The node colors represent different network objects (histone PTMs, metabolites, and drugs), while the node size represents the absolute value of the object. We used a p-value < 0.05 and FDR < 0.25 from the Pearson Correlation Coefficients to filter out non-significant interactions. We used a liberal statistical threshold since we combined multiple methods to identify the most robust interactions. For LASSO networks with many nodes, we prioritized interactions that were the strongest in their respective datasets by selecting the upper and lower quartiles of the LASSO coefficients.

### Experimental metabolite and histone PTM inhibitor sensitivity profiling

We evaluated drug/metabolite synergism and antagonism on A549 cell lines (ATCC), using our model predictions and correlation analysis as a guide to select nutrient supplementation conditions and histone PTM inhibitors. Using an automated platform, A549 was plated at a density of 200 cells per well in 384 well plates. Cells were grown in their respective media using standard cell culture conditions (37°C, 5%). The effect of small molecules was measured over an 8-point concentration range with three concentrations of metabolites (195, 625, and 2000 uM) with < 0.5% DMSO. Media was supplemented with D-Sorbitol (MilliporeSigma S1876-100G), 1-methylnicotinamide (Fisher Scientific 11-101-2274), reduced glutathione-ester (MilliporeSigma G6013-5G), kynurenic acid (MilliporeSigma K3375-250MG), DL homocysteine (MilliporeSigma H4628-1G), 5-adenosylhomocysteine (MilliporeSigma A9384-100MG), and D-pantothenic acid (MilliporeSigma P5155-100G). Cells were treated with histone deacetylase inhibitors Vorinostat (MilliporeSigma SML0061-5MG) and histone demethylase inhibitor GSK-J4 (MilliporeSigma SML0701-5MG). We assessed cell count over 48 hours after compound/drug pairwise treatment as a surrogate for cell viability. For studying the impact of drugs and metabolites on histone PTMs, total histones were isolated by using EpiQuik Total Histone Extraction Kit (Epigentek, cat. No OP-0006), according to manufacturer instructions. Quantification of 21 histone H3 modifications was performed by using the EpiQuik Histone H3 Modification Multiplex Assay Kit (Epigentek cat no. P-3100), according to manufacturer instructions. For the follow up screen to assess interactions at lower metabolite doses, we used the same experimental setup to confirm synergistic interaction between the drugs and kynurenic acid, 1-MNAM, and pantothenate. These metabolites were prioritized by our models as they showed synergistic interaction with either of the drugs experimentally (**see Table 1**).

### Drug sensitivity data processing

Cell count data (N=2 treatment replicates; N=12 control replicates) was normalized using the average DMSO and drug-only cell counts at each compound and drug concentration. Dose-response curves for each metabolite/drug pair were generated using the DMSO and drug-only normalized cell count data. To determine drug/metabolite synergy and antagonism, we calculated the cell count percentage in the drug/metabolite treatment over the drug-only condition. Each metabolite/drug pair was visualized as a dose-response matrix to visualize synergism or antagonism. We calculated the Bliss scores for each of the combinations using the following formula^101^:

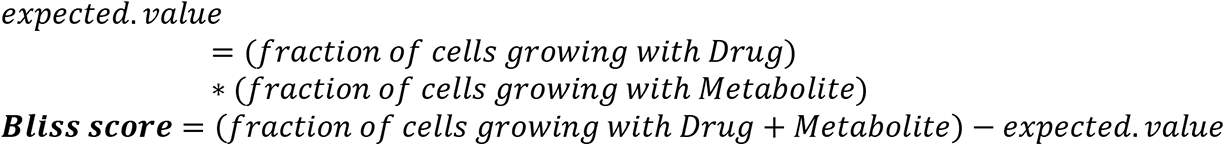

Cell counts with DMSO were taken as the control. All data points were averaged across two replicates. Negative bliss scores represent synergistic interactions and position scores imply antagonism.

## Acknowledgments

This work was supported by faculty start-up funds from the University of Michigan (UM), Camille and Henry Dreyfus Foundation, the UM Rogel Cancer Center, UM Endowment for Basic Sciences Accelerator Award, UM Research Scouts Award, the UM Chad Carr Pediatric Brain Tumor Center, and R35 GM13779501 from NIH to SC, and the Proteogenomics of Cancer Training Program to SEC. We thank Peter Toogood and Andy Alt at the Michigan Drug Discovery core for discussions on the experimental assays, Claudia Lalancette for assistance with the epigenetics experiments, Costas Lyssiotis, Yali Dou, Tim Cernak, Sriram Venneti, Duxin Sun, and Neil Youngson for suggestions and critical feedback on the project.

## Data availability

All datasets are provided in figures and in the supplementary tables.

## Author contributions

S.C conceived the study, designed, and performed research, S.E.C, R.B, T.L, A.S, R.J, A.R, M.S, designed and performed research, and S.C, R.B and S.E.C wrote the manuscript.

## Supplementary Figures

**Supplementary Figure 1** Correlation profiles for metabolite/PTM correlations using both metabolite and lipid data

**Supplementary Figure 2** Correlation profile between histone PTMs and drug sensitivity

**Supplementary Figure 3** LASSO metabolomics/histone PTM CCLE predictions

**Supplementary Figure 4** LASSO drug - metabolite interactions

**Supplementary Figure 5** Stepwise regression predictions for metabolite/PTM and metabolite/AUC interactions

**Supplementary Figure 6** 1-MNAM and H3K9me3 k-TSP predictions

**Supplementary Figure 7** Assessing the metabolic impact on histone PTM status during EMT

**Supplementary Figure 8** Metabolite-epigenetic drug dose-response curves for A549 using DMSO control

**Supplementary Figure 9** Tile plots of bliss scores calculated for drug - metabolite combinations in A549 cells

**Supplementary Figure10** Kynurenic acid-GSK-J4 synergistic interaction and dose-response curves for HCT116

**Supplementary Figure 11** Metabolite/drug 2D dose-response profiles for Vorinostat and GSK-J4 in A549

## Supplementary Tables: Supplementary_Tables.xlsx

**S. Table 1** Pearson correlation coefficients between metabolite/PTM interactions

**S. Table 2** Literature validation of metabolite/histone PTM interactions

**S. Table 3** Pearson correlation coefficients between metabolites and drug sensitivity data

**S. Table 4** Pearson correlation coefficients between drug sensitivity data and histone PTM levels

**S. Table 5** LASSO Met/PTM interactions

**S. Table 6** LASSO Met/AUC interactions

**S. Table 7** Stepwise regression interactions

**S. Table 8** Aggregated ML metabolite/PTM model (top 20 metabolites)

**S. Table 9** Aggregated ML metabolite/Drug model (top 20 metabolites)

**S. Table 10** Summary of metabolite/PTM interactions predicted by correlations and ML

**S. Table 11** Summary of metabolite/drug interactions predicted by correlations and ML

**S. Table 12** t-statistics of linear regression model between genetic mutations and metabolite abundance.

**S. Table 13** t-statistics of linear regression model between copy number variation and metabolite abundance.

**S. Table 14** Correlation between Drug sensitivity and mutations across DepMap cell lines

